# hAMRonization: Enhancing antimicrobial resistance prediction using the PHA4GE AMR detection specification and tooling

**DOI:** 10.1101/2024.03.07.583950

**Authors:** Inês Mendes, Emma Griffiths, Alex Manuele, Dan Fornika, Simon H Tausch, Thanh Le-Viet, Jody Phelan, Conor J. Meehan, Amogelang R. Raphenya, Brian Alcock, Elizabeth Culp, Federico Lorenzo, Maria Sol Haim, Adam Witney, Allison Black, Lee Katz, Paul Oluniyi, Idowu Olawoye, Ruth Timme, Hui-min Neoh, Su Datt Lam, Tengku Zetty Maztura Tengku Jamaluddin, Sheila Nathan, Mia Yang Ang, Sabrina Di Gregorio, Koen Vandelannoote, Rutaiwan Dusadeepong, Leonid Chindelevitch, Muhammad Ibtisam Nasar, David Aanensen, Ayorinde Oluwatobiloba Afolayan, Erkison Ewomazino Odih, Andrew Grant McArthur, Michael Feldgarden, Marcelo M Galas, Josefina Campos, Iruka N. Okeke, Anthony Underwood, Andrew J. Page, Duncan MacCannell, Finlay Maguire

## Abstract

The detection of antimicrobial resistance (AMR) markers directly from genomic or metagenomic data is becoming a standard clinical and public health procedure. This has resulted in the development of a number of different bioinformatic AMR prediction tools. Although many may implement similar principles, these tools differ significantly in their supported inputs, search algorithms, parameterisation, and underlying reference databases. Each of these tools generates a report of detected AMR genes or variants in a distinct, non-standard, format. This presents a huge barrier to the comparison of results and to the modularity of tools for AMR gene prediction within bioinformatic workflows.

In collaboration with 17 public health laboratories across 10 countries, the Public Health Alliance for Genomic Epidemiology (PHA4GE) (https://pha4ge.org) data structures working group has developed and piloted a standardized output specification for the bioinformatic detection of AMR from microbial genomes. In this report, we discuss hAMRonization, a python package and command-line utility, which implements PHA4GE’s AMR specification to combine the outputs of disparate antimicrobial resistance gene detection tools into a single unified format. hAMRonization can be easily extended and currently supports 18 different tools (both species-agnostic and species-specific) for the detection of genes and/or variants conferring AMR. The harmonized reports are available in tabular form, JSON format or through an interactive HTML file (e.g., https://maguire-lab.github.io/assets/interactive_report_demo.html) that can be opened within the browser for navigable data exploration. As of 2024-03-07 hAMRonization has been downloaded ∼12,500 times, incorporated into >9 public bioinformatic tools and workflows, and been internally adopted by several national and international public health groups. The hAMRonization tool and underlying specification are open-source and freely available through PyPI, conda and GitHub (https://github.com/pha4ge/hAMRonization).

## Introduction

Antimicrobial resistance (AMR) represents a growing public health crisis with extensive global impact across human and animal health, agriculture, and trade. Multidrug resistance is increasing in a broad range of bacterial pathogens [1]; this, combined with low rates of novel antimicrobial drug discovery and development [2], underscores the importance of addressing this threat [3]. National and international antimicrobial resistance action plans e.g., [3–6], have identified several strategies to mitigate the risk of AMR, such as rapid detection of the AMR determinants present within a clinical sample, improved surveillance of AMR, more targeted antimicrobial use, applied research into resistance mechanisms, and a better understanding of the mechanisms driving environmental transmission of AMR determinants.

Diagnostic and public health surveillance are increasingly using accredited analyses of genomic and metagenomic data for pathogen identification and characterization [7,8]. Therefore, accurate identification of genes or variants associated with potential AMR from genomic data is critical for detecting, monitoring and attempting to mitigate the spread of AMR. Genomic surveillance and outbreak investigations for resistant pathogens are already being performed all over the world by public health agencies, clinicians, industry, and academic researchers, often without much standardization. Given the scale of this problem and the number of stakeholders involved, many bioinformatics tools have been developed that are dedicated to the task of AMR gene and variant detection [7,9,10]. As of March 2024, there are more than 18 active open-source, command-line tools designed to identify the presence of genes and/or variants associated with AMR, and a number of other proprietary solutions that include AMR genotyping data [9].

These tools include several developed to work with a specific primary species-agnostic database, e.g., the Resistance Gene Identifier (RGI) for the Comprehensive Antibiotic Resistance Database (CARD) [11], AMRFinderPlus and the National Center for Biotechnology Information (NCBI) Pathogen Detection Reference Gene catalog [12,13], and ResFinder [14] and KmerResistance [15] for the ResFinder database [14]. Other tools exist that use merged forms of existing databases such as ResFams [16], AMRplusplus [17], and DeepARG [18]. There are also organism-specific tools designed to handle the specific and nuanced challenges of certain taxa such as Kleboroate for *Klebsiella pneumoniae* species complex [19] or TB-Profiler for *Mycobacterium tuberculosis* [20]. Finally, there are AMR gene identification tools that provide a database-agnostic approach using novel algorithms (e.g., GROOT [21], ARIBA [22], Mykrobe [23]), or convenient interfaces (e.g., ABRicate [24], sraX [25]).

These tools exhibit different strengths and weaknesses due to differences in underlying databases, search algorithms, and default parameterisations. Based on the specific requirements of a given AMR analysis some tools may be better suited to the analytical workflow than others. Additionally, each of these tools generates differently formatted outputs and reports, all in non-standard formats, and all using inconsistent terminology and interpretive criteria. This poses a significant challenge to effective integration, modularity, and comparison of AMR gene and variant detection methods. The consequence of this is that it is difficult for researchers and public health experts to systematically evaluate the suitability of different tools in their workflow. The limited examples of these comparisons are largely reliant on the development of custom ad-hoc tool-specific parsers [12,22], or have skipped the tools entirely and directly compared the underlying search algorithms [26]. Even in cases where benchmarking has been performed, the disparate output formats mean it requires significant work to modify a workflow to use a different AMR detection tool. This greatly limits modularity and flexibility to integrate new tools into existing analyses or repurpose them to new or changing requirements. Given the recent calls to mitigate these issues in public health genomic epidemiology [27], it is critical that data structures and methods are developed which enable tool-agnostic, robust parsing, manipulation, and transformation of AMR gene detection results.

To this end, via an iterative expert-led development and pilot process across 17 public health laboratories in 10 countries, we compared and consolidated the outputs of existing AMR gene detection tools to develop the hAMRonization specification. This is a standardized set of recommended and mandatory output terms and labels for AMR gene detection tools, such as “Gene Name”, “% Coverage (breadth)”, and “Drug Class”. The outputs for all 18 currently maintained, species agnostic, open-source AMR gene detection tools can be directly converted to this unified specification. Furthermore, this conversion to a common AMR gene detection output specification can be performed automatically using a companion tool and library of biopython compatible parsers.

Through the uptake of the hAMRonization data standard and associated tools, we aim to improve the comparability of results, and the interoperability of workflows in the public health microbial genomics ecosystem. This will lead to better AMR surveillance, more effective genomic diagnostics, and ultimately improved human and animal health.

## Materials and Methods

### The PHA4GE hAMRonization Specification

To compare and unify the broad set of output fields across individual AMR gene detection tools it is necessary to create a common language that can be used to describe them all. This takes the form of a standardized set of output labels and ontological identifiers (linked to standardized definitions) (Figure 1). Individual fields from the output of each tool can then be mapped to a standardized label. For example, the accession of the contig a given AMR gene was detected on is listed as “Contig” in RGI, “contig_name” in ResFinder, and ‘contig id’ in AMRFinderPlus. Therefore, our specification includes a standardized “Contig ID” field which captures all of these values across tools.

**Figure 1:**
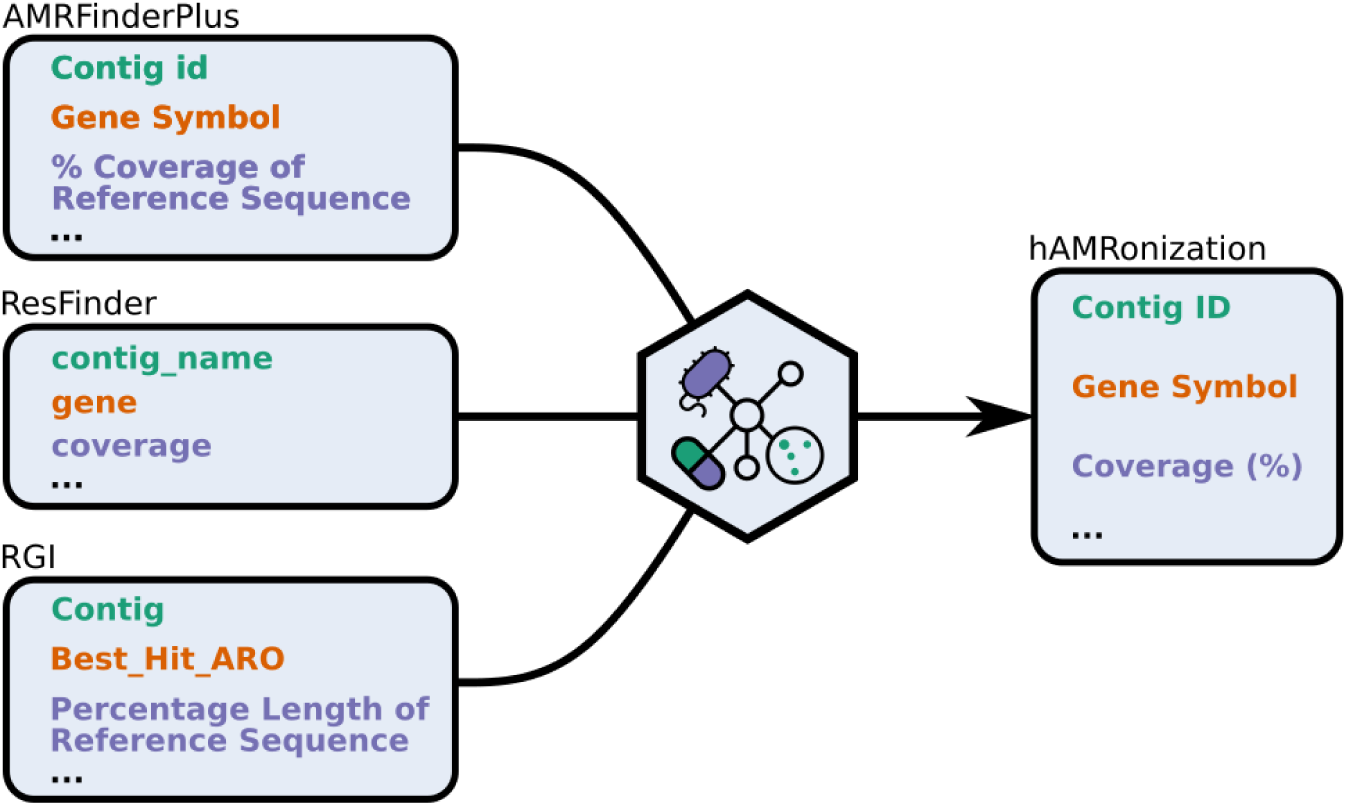
The PHA4GE hAMRonization specification schematic. The curating and comparing output formats from 18 open-source AMR gene detection tools, with evidence of active use in the literature, was performed and summarized in a core set of essential contextual data for use of AMR detection in health or research contexts. The figure illustrates several output categories from AMRFinderPlus, ResFinder and RGI that were harmonized into three distinct terms.

This common language, or specification, was created through an iterative expert-led process which consisted of curating and comparing output formats from 18 open-source AMR gene detection tools (see Supplemental Table 2), with evidence of active use in the literature. This included consultation with an international set of national public health groups, tool developers, and AMR genomics experts through the PHA4GE initiative. In addition to identifying terms compatible with the range of outputs, this work identified a core set of essential contextual data for use in AMR detection in health or research contexts. This includes items such as the input filename, name and version of the AMR database used, and the name and version of the AMR gene detection tool. In total, the specification consists of 35 fields that take into account the variability in the way information is expressed across tools (e.g. breadth of coverage expressed as a percentage (90%) vs a ratio (450/500)). Once the prioritized data elements were selected, definitions and implementation guidance for the standardized fields were developed. Appropriate ontology terms matching the desired data elements were sourced from different OBO Foundry (https://obofoundry.org/) [28] ontologies. Ontologies within the OBO Foundry library are developed using common principles and practices designed to increase interoperability across domains of knowledge [29]. Terms in the draft AMR specification for which ontology concepts already existed were adjusted to align with community standards. Terms which did not already exist were submitted to appropriate source ontologies. Ontologies are meant to represent “universal truth” as much as possible, and not just use case-specific nuances. As such, additional guidance was provided for AMR detection-specific implementation. Fields, definitions, specification-specific guidance, value types, and examples of use can be found in the hAMRonization specification available in GitHub (https://github.com/pha4ge/hAMRonization/tree/master/schema), encoded both in a human-readable CSV file and a JSON format to enable automated computer validation. Standardized vocabulary was sourced/harmonized with the Ontology for Biological Investigations (OBI) [30], the Antimicrobial Resistance Ontology (ARO) [11], the Genomic Epidemiology Ontology (GenEpiO) [31], and the Chemical Entities of Biological Interest Ontology (ChEBI) [32].

### The hAMRonization Package

Mappings from the output fields in each of the 18 tools to the hAMRonization specification were encoded in a python package (https://github.com/pha4ge/hAMRonization). This package was developed following the extensible modular approach of biopython [33] and provides an easy-to-use command line tool to enable the automatic conversion of output files from each of these tools to a standardized report following the hAMRonization specification. Where appropriate, and when tools do not provide the full set of essential core metadata, hAMRonization will prompt the user for this information to ensure high-quality reproducible analyses. This tool also supports the collation of all the hAMRonized results from multiple tools and samples into a single report to aid comparison and analysis. These reports can optionally be generated as spreadsheets (CSV), JSON, or an interactive navigable HTML format to maximize utility and compatibility with downstream processes. Finally, this package provides a python module to enable direct integration of these functions into other python-based tools. To ensure practical installation in as broad a range of computing infrastructures as are present across public health, academic, and industry research settings this tool is open source and is freely available via GitHub, PyPI, bioconda, docker, and the galaxy toolshed.

### The hAMRonization Proof of Concept Workflow

As a proof-of-concept, a snakemake pipeline [34] was created which automatically runs arbitrary microbial genomic data through a set of species-agnostic AMR gene detection tools, supported by hAMRonization, and generates a single report for each. This set is composed of outputs from (a subset of supported tools): abricate (https://github.com/tseemann/abricate, version 1.0.1), amrfinderplus [12] (version 3.10.1), amrplusplus [17] (v1.0.0), ariba [22] (version 2.14.6), c-SSTAR [35] (version 1.1.01), groot [21] (version 1.1.2), kmerresistance [15] (version 2.2.0), resfams [16] via hmmer (v3.3.2), resfinder [14] (version 3.2), rgi [36] (version 5.1.1), srax [25] (version 1.5), srst2 [37] (version 0.2.0) and staramr [38] (version 0.7.2). In order to guarantee the reproducibility of the results obtained, it integrates fixed versions of the tools implemented in the pipeline from conda. The pipeline is available at https://github.com/pha4ge/hAMRonization_workflow (version 1.0.0).

## Results

The PHA4GE hAMRonization Detection specification was designed to harmonize information prioritized by public health practitioners for genomic surveillance, outbreak investigations and research with direct public health impacts. Fields from various widely used AMR detection tools were compared, and a set of 35 fields were selected to capture information regarding the software and database provenance, information about reference genes and genomes, as well as mutations and resistance genes detected in sequences of interest (Supplemental Table 1). The field labels are standardized using terms sourced or contributed to open-source ontologies. Ontologies are sets of controlled vocabulary arranged in a hierarchy, in which terms are linked using logical relationships and the meanings of the terms are disambiguated by the assignment of unique and persistent identifiers. The positions of detected mutations with respect to reference genes/genomes/proteins are specified using the machine-readable Human Genome Variation Society (HGVS) notation system, a sequence variant nomenclature that has been used previously for encoding *Mycobacterium tuberculosis* drug resistance mutations [39,40].

### hAMRonization Data Transformation, Parsing Tools, and Harmonized Reports

To enable automated data transformations from variable tool outputs to the PHA4GE hAMRonization specification, fields from 18 gene and mutation detection tools (i.e., ABRicate [24], AMRFinderPlus [13], AMRPlusPlus [17], ARIBA [22], C-SSTAR [35], DeepARG [18], fARGene [41], GROOT [21], KmerResistance [15], Mykrobe [23], PointFinder [42], ResFams [16], ResFinder [14], RGI [11], SraX [25], SRST2 [37], StarAMR [38], and TBProfiler [20]) were mapped to the specification (Figure 2). It is worth noting that not all tools provide values for all specification fields. Also, many tools provide outputs beyond what is captured by the specification (e.g. “Model_type” in RGI, “Haplotype pattern” in AMRPlusPlus). These values are excluded from the harmonized report.

**Figure 2:**
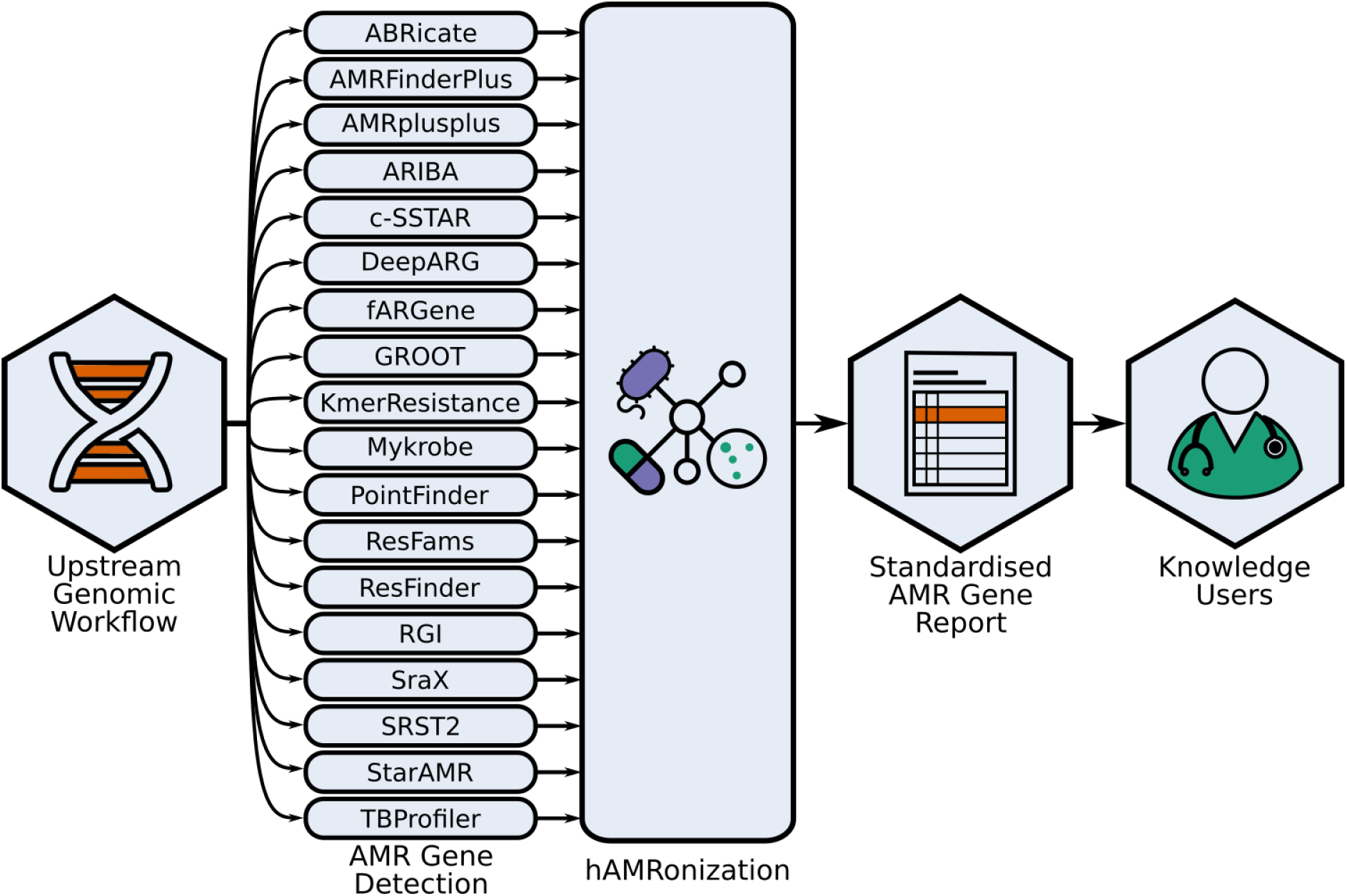
Overview of hAMRonization within standard AMR genomics/metagenomics workflows. The hAMRonization module and command-line utility tools allow the creation of a single unified output report from any of 18 established disparate AMR gene and/or variant prediction tools. This enables users such as national public health reference laboratories, clinical laboratories, and academic groups to compare results across tools and change their workflow to different tools without having to develop custom code. It also means that the communication of results to downstream knowledge users, such as infection prevention and control teams, can be done in a consistent standardized manner regardless of which AMR prediction tool was used in the genomic/metagenomic analysis.

Parsers were then constructed to automate the transformation and movement of outputs into the appropriate fields in the hAMRonization reports. Parser code is open source, and all code and installation instructions are available at https://github.com/pha4ge/hAMRonization. hAMRonization can be executed from the command-line, and alternatively, can be installed and used in Galaxy via the Galaxy tool shed. hAMRonization provides both tabular and interactive reporting formats. The interactive report provides the user with an HTML file that can be opened within the browser for navigable data exploration.

### Piloting hAMRonization in Real-World Public Health Settings

To test the utility and value of the hAMRonization specification and its associated tools for AMR genomic surveillance and data sharing, pilot projects were launched in laboratories across three partner teams, which included Cambodia/Australia, Nigeria (seven laboratories across the AMR surveillance network), and Malaysia/Argentina via a PHA4GE subgrant competition (Figure 3). PHA4GE also engaged eight labs from seven countries in Latin America via the ReLAVRA AMR surveillance network (a network associated with the Pan American Health Organization (PAHO), a regional office of the World Health Organization). All pilot project participants were issued with a short document outlining the purpose of the software, installation instructions and prompting questions for feedback on their experience. Requested feedback focused on the ease of use of the tool, as well as its utility in sharing harmonized data among their networks where previously sharing was difficult due to the variability of tool outputs.

**Figure 3:**
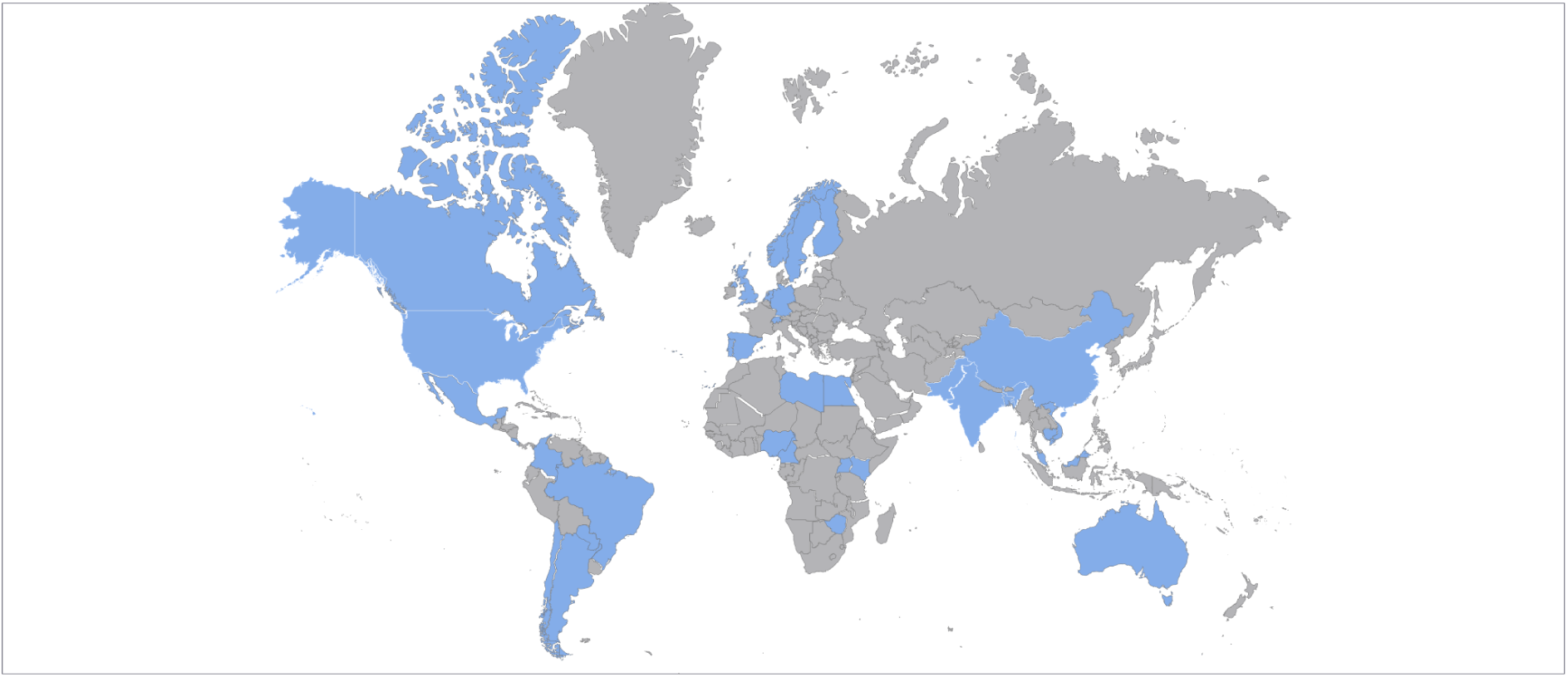
Countries where hAMRonization was piloted. Pilot projects were launched in laboratories across three partner teams, which included Cambodia/Australia, Nigeria (seven laboratories across the AMR surveillance network), and Malaysia/Argentina via a PHA4GE subgrant competition. PHA4GE also engaged eight labs from seven countries in Latin America via the ReLAVRA AMR surveillance network.

Subgrant participants used hAMRonization to analyze data from hospital settings and/or national surveillance projects involving a variety of organisms e.g. *Salmonella enterica* ser. Paratyphi A [43] and Methicillin-Resistant *Staphylococcus aureus* (MRSA). Feedback included the need for improved definitions of fields and guidance on required inputs, the need for identifying causes of hAMRonization crashes, and the need for more clinician-friendly installation and execution instructions. Some participants also suggested that the inclusion of sequence contextual data would be helpful for analysis (i.e., hospital name and country of sample collection, ward (medical vs surgical), collection date, specimen type (e.g. blood, urine), latitude/longitude of sample collection). In addition to the testing exercise performed, the Nigeria Team (led by the University of Ibadan) integrated hAMRonization into their NextflowTower platform for AMR bioinformatics analysis training workshops. To better enable their clinician colleagues without bioinformatics expertise to use the hAMRonization tool, the Malaysia/Argentina Team (led by the National University of Malaysia) developed a Google Colaboratory (a Google Notebook that enables code to be executed from a Browser) and has distributed it to their network partners. This has also included running 2 virtual training workshops attended by clinicians, veterinarians, and public health professionals from Malaysia, Argentina, and Japan.

The ReLAVRA pilot included National Reference Laboratories from Argentina, Brasil, Chile, Colombia, Costa Rica, México and Paraguay. These laboratories were provided with a dataset of 20 *Shigella sonnei* genomes from a previous Latin America regional project analyzed with 5 different AMR detection tools: ABRIcate, AMRFinderPlus, ARIBA, SRST2, StarAMR. Participants received the outputs of the AMR detection tools together with the instructions for installing and running hAMRonization. As hAMRonization and its associated materials are written in English, the documentation had to first be translated by PAHO prior to distribution in Spanish-speaking Latin American countries. The pilot instructions (in both English and Spanish) are available at https://github.com/pha4ge/hAMRonization/tree/master/docs/subgrant. When participants completed the exercise, they were asked to complete a structured poll. Twelve responses from the seven participating nations were received, and a summary report was prepared by ReLAVRA and provided to PHA4GE. Most of the participants were bioinformaticians and laboratory technicians. For bioinformaticians, the installation instructions were easy to follow, however, laboratory technicians without a bioinformatics background expressed difficulties largely with the dependency installation - a barrier that could be overcome in the future by making all tools available as Bioconda packages. Participants agreed that all harmonized output fields provided relevant information for AMR detection but the most useful were the fields that cover the AMR gene detected and the confidence in that result. Respondents were concerned about the importance of having knowledge of AMR and genomic data analysis for the correct interpretation of the results, especially with regard to decision-makers in clinical settings. This challenge is a barrier for the implementation of genomics for AMR surveillance in clinical settings and should be addressed. Unfortunately, clinical interpretation and phenotype prediction from genomic data in general is a major open problem [14,44–46] that requires careful contextualisation of identified resistance genes in their clinical (infection type, role of biofilms, quorum sensing and so on), microbial (underlying organismal biology and other genes), bioinformatic (what types of resistance can a specific tool detect e.g., internal stop codons), and functional (what is the specific mechanism of this gene) aspects. Organism-specific tools (e.g., TB-profiler) can aid clinical interpretation but being more optimized around these nuances and using hAMRonization can make it easier to use these without having to design custom workflows around them.

In terms of output summary format, most of the participants preferred the tabular report, indicating that a single file combining all the relevant information was more convenient and amenable to processing with different downstream software. However, some participants regarded the interactive HTML report as helpful for comparing which tools identified the same gene.

Overall, all users reported finding the tool very useful for regional genomic surveillance and sharing results with different laboratories across networks employing a range of AMR detection tools. Technical issues such as fixes for crashing, improved definitions, and instructions for inputs were addressed. Ways to address bigger picture requests such as the harmonization of gene nomenclature, tool-specific issues, and the inclusion of contextual data (sample metadata as well as epidemiological, clinical and laboratory data) are a natural extension of this work and actively under discussion within PHA4GE.

### hAMRonization and Hackathon-based Community Development

A hackathon jointly organized by PHA4GE, CLIMB-Big Data, and JPIAMR was held in October 2021 (https://github.com/AMR-Hackathon-2021). The aim of the hackathon was to improve upon currently available bioinformatics tools and methods for the AMR community, with a special focus on antimicrobial resistance in bacteria. A hackathon is an event in which a large number of people from the community meet to engage in collaborative computer programming. Project ideas are pitched, and participants form working groups based on the projects proposed. PHA4GE representatives pitched challenges identified during the hAMRonization pilot implementations as hackathon projects - such as the need for a clinician-friendly interpretation of HGVS mutation notation and the need for harmonization of gene nomenclature across reference databases. As a result of the collaborative work during the hackathon, the hAMRonization specification was improved by the development of a layman’s translation of HGVS notation specifying the coordinates of mutations identified within genes, proteins and genomic sequences of interest in relation to their position in a reference sequence (https://github.com/conmeehan/laymansHGVS). Furthermore, the use of an ontology-based tool for harmonizing gene nomenclature called CHAMREdb (https://gitlab.com/antunderwood/chamredb) was explored and tested in hARMonization. While CHAMREdb is not yet fully integrated into hAMRonization, we anticipate its involvement in future development. These achievements are examples of the important work that can be done when scientists are brought together to use their creativity and expertise to solve problems at community-driven hackathons.

### Further adoption and use of hAMRonization

Beyond the adoption described above, hAMRonization has also seen adoption from the broader community in the form of “best practices” guides (e.g., [47–49]), bioinformatics tools/libraries (e.g., nf-core module [50], funcscan (https://github.com/nf-core/funcscan), graphAMR [51], hAMRoaster [52], argNorm (https://github.com/BigDataBiology/argNorm)), production public health workflows (e.g., Latvia’s ARDETYPE workflow (https://github.com/NMRL/Ardetype) and Norway’s metagenonix workflows (https://github.com/FranckLejzerowicz/metagenomix)), and research papers (e.g., [53–57]). As of 2024-03-07, hAMRonization has been download ∼12,500 times from PyPI, galaxy toolshed, and bioconda repositories.

## Discussion

The lack of standardization in the reporting of AMR gene and mutation detection greatly hinders the comparison of results across the public health sector. The myriad of options available for this purpose highlights a severe interoperability problem. In this work, we have identified and prioritized data elements capturing AMR detection information most pertinent for public health decision-making. We have developed a standardized specification for harmonizing variable AMR detection tool outputs, implementing open source, community-based ontologies as well as standardized formats and nomenclatures. We have also operationalized the specification through the development of tools (parsers and workflows) and harmonized reports, available as a package called hAMRonization. Finally, we have tested the hAMRonization in different public health and research settings. The results of this work show that hARMonization enables the dissemination of results to stakeholders in a single consistent format, allowing not only the comparison of tools and databases but the validation of results through multiple detection algorithms.

The developed standardized data specification and tooling improve data harmonization and interoperability, allowing the comparison of results not only across multiple AMR detection algorithms but also AMR reference databases. hAMRonization also improves the capture of vital data provenance information, such as the versions of software and databases used in analyses, which are necessary for reproducibility and, increasingly, for accreditation processes [8].

It should be noted that while hAMRonization has been applied to AMR detection, many data elements are case-agnostic and so can be repurposed for the detection of genes and mutations associated with other phenotypes such as virulence, pathogenicity, transmissibility, heavy metal resistance, mobility factors etc. Extension of the specification through the adding of other optional higher-order annotation fields such as virulence factor type (“adherence”, “toxin” etc) or metal type, could help to create modularity, different types of detection streams, and feed directly into other downstream analysis. A limitation of the current hAMRonization specification is that while individual determinants contributing to resistance can be detected, there is no support for highlighting the presence of multi-component resistance mechanisms such as glycopeptide resistance clusters [58] or efflux pumps [59].

One of the most substantial challenges with any attempt to improve interoperability by introducing new standards/specifications is their uptake. We have attempted to minimize the barriers to uptake by operationalizing this specification with a set of automated data transformation tools. For each of the AMR detection tools, a set of “mappings” or instructions to convert their outputs to the hAMRonization specification has been provided along with parsers to automate this process. Ideally, developers will adopt this specification and structure their tool’s output accordingly in the future. This would obviate the need for future mappings and parsers. However, even if this does not occur, the nature of the specification means that only minimal effort is required to convert tabular results from any novel AMR gene detection tool. Another benefit of such a community-based consensus standard for AMR detection is that it provides much-needed standards for industrial partners developing instruments and devices. A challenge within public health laboratories is the lack of interoperability between different phenotype characterization instruments and lab information management systems. Providing community-vetted, standardized specifications for genomics applications enables vendors to develop tools and other resources that are interoperable and add value (for example, an AMR typing platform conforming to the hAMRonization standard could produce results that could more easily plug into a LIMS or other downstream software more seamlessly).

The implementation of hAMRonization in real-world settings has enabled us to clearly catalog and articulate public health challenges outside of the scope of the specification and tools, that impact the use of hAMRonization. Lessons learned include the fact that the harmonization of data for genomic surveillance comes with additional needs outside of standards and technical implementation, including improved support for the interpretation and evaluation of harmonized data. Also, in harmonizing the outputs of tools using different computational approaches, the strengths and weaknesses of the tools need to be made more transparent to users as they become removed from working directly with the individual tools themselves. Future work will focus on these areas, as well as better harmonizing gene nomenclature across reference databases.

In summary, through the uptake of the hAMRonization data standard and associated tools, we aim to improve the comparability of results, and the interoperability of workflows in the public health microbial genomics ecosystem. This will lead to better AMR surveillance, more effective genomic diagnostics, and ultimately improved human and animal health.

## Acknowledgements

The authors would like to thank all contributors to hAMRonization on GitHub. Additionally, the authors acknowledge all participants of the AMR hackathon organized by PHA4GE, CLIMB-Big Data, and the Joint Programming Initiative on Antimicrobial Resistance (JPIAMR), and the Latin American Public Health Laboratories via the ReLAVRA AMR surveillance network that participated in the pilot test. The findings and conclusions in this report are those of the author(s) and do not necessarily represent the official position of the Centers for Disease Control and Prevention (USA), National Institutes of Health (USA), or any other government agency affiliated with an author. The work included was reviewed by the Centers for Disease Control and Prevention (USA) and was conducted consistent with applicable federal law and Centers for Disease Control and Prevention policy (e.g., 45 C.F.R. part 46, 21 C.F.R. part 56, 42 U.S.C; 241(d); 5.U.S.C §552a; 44 U.S.C. §3501 et seq).

## Disclaimer

Nothing to declare.

---

**Supplemental Table 1:** hAMRonization: AMR Gene Detection Output Specification.

| Interface Label | Required/Optional | Definition | Ontology Identifier | Value Type | Example | Guidance | Values |
| --- | --- | --- | --- | --- | --- | --- | --- |
| **Analysis Software Name** | Required | A name of a computer package, application, method or function used for the analysis of data. | OBI:0002894 | String | amrfinder | Not typically included in an AMR prediction output. The user may have to provide the value. |  |
| **Analysis Software Version** | Required | The version of software used to analyze data. | OBI:0002896 | String | 1.2.5 | The version of the software revealed by the version flag such as "rgi main --version" or "abricate --version". Not typically included in an AMR prediction output. Semantic versioning is strongly recommended. If unavailable, provide the date the tool was last updated or installed in ISO 8601 format (yyyy-mm-dd). |  |
| **Gene Name** | Required | The name of a gene, (typically) assigned by a person and/or according to a naming scheme. It may contain white space characters and is typically more intuitive and readable than a gene symbol. It (typically) may be used to identify similar genes in different species and to derive a gene symbol. | OBI:0002878 | String | type A-1<br>chloramphenicol<br>O-acetyltransferase | The long name for the gene/protein. Gene names or gene product names are acceptable. |  |
| **Gene Symbol** | Required | The short name of a gene; a single word that does not contain white space characters. It is typically derived from the gene name. | OBI:0002877 | String | catA1, blaOXA-101 | Acceptable values may represent gene symbols or gene product symbols. "Gene symbol" and "allele" (based on amino acid variations) may be used interchangeably due to nuances in resistance gene family nomenclature. Parsers should include values from the "gene symbol" and "allele" fields in this category. |  |
| **Genetic Variation Type** | Required | The class of genetic variation detected. | GENEPIO:0100401 | Enums | Gene detected | Select a value from the pick list. | ['Gene presence detected', |
|  |  |  |  |  |  |  | 'Mutation detected', 'Both gene presence and mutation detected'] |
| **Input File Name** | Required | The name of the file containing the sequence data to be analysed. | OBI:0002874 | String | PDT000607292_1_genomic.fna | Name of the file with the input sequence data to be analysed. Tools can have multiple input files (nucleotide space/amino acid space and/ or paired-end read files), in which case they should be separated by a semi-colon delimiter. Multiple input files will be accommodated by their corresponding parser. |  |
| **Reference Accession** | Required | An identifier that specifies an individual sequence record in a public sequence repository. | OBI:0002885 | String | NF000491.1 | No specific guidance. |  |
| **Reference Database Name** | Required | An identifier of a biological or bioinformatics database. | OBI:0002883 | String | NCBI, ResFinder | The name of the reference database may need to be entered manually if not automatically recorded as a parameter within the tool. |  |
| **Reference Database Version** | Required | The version of the database containing the reference sequences used for analysis. | OBI:0002884 | String | 1.2.3 | The version of the reference database may need to be entered manually if not automatically recorded as a parameter within the tool. If unavailable, provide the date the database was last updated or installed in ISO 8601 format (yyyy-mm-dd). |  |
| **Antimicrobial Agent** | Optional | A substance that kills or slows the growth of microorganisms, including bacteria, viruses, fungi and protozoans. | CHEBI:33281 | String | CHLORAMPHENICOL | This should describe a specific agent. Standardized terms from the ChEBI ontology should be used. Find terms using https://www.ebi.ac.uk/ols/ontologies/chebi. Can be left blank if not provided. |  |
| **Coverage (percentage)** | Optional | The percentage of the reference sequence covered by the sequence of interest. | OBI:0002880 | Float | 90 | The sequences can refer to reads and genome, nucleotides and genes, or amino acids and proteins. This value should be normalized and expressed as a percentage. Do not include the "%" sign in the value. |  |
| **Coverage (depth)** | Optional | The average number of reads representing a given nucleotide in the reconstructed sequence. | GENEPIO:0000092 | Float | 56 | The value is expressed as a fold value e.g. 56x. |  |
| **Coverage (ratio)** | Optional | The ratio of the reference sequence covered by the sequence of interest. | OBI:0002881 | String | 450/500 | The sequences can refer to reads and genome, nucleotides and genes, or amino acids and proteins. This value should not be normalized and expressed as the ratio of actual positions being compared e.g. 450/500. |  |
| **Drug Class** | Optional | In an antimicrobial context, a drug class is a set of antibiotic molecules, including antibiotic/adjuvant combination medications, with similar chemical structures, molecular targets, and/or modes and mechanisms of action. | ARO:3005165 | String | Phenicol | Standardized terms from the ChEBI ontology should be used. Find terms using https://www.ebi.ac.uk/ols/ontologies/chebi. Can be left blank if not provided. |  |
| **Input Gene Length** | Optional | The length (number of positions) of a target gene sequence submitted by a user. | OBI:0002891 | Int | 657 | Report the number of nucleotides in the sequence without any units. |  |
| **Input Gene Start** | Optional | The position of the first nucleotide in a gene sequence being analyzed (input gene sequence). | OBI:0002977 | Int | 18 | The value should be biologically relevant, and should not start with 0. Can be left blank if not provided. |  |
| **Input Gene Stop** | Optional | The position of the last nucleotide in a gene sequence being analyzed (input gene sequence). | OBI:0002978 | Int | 921 | The value should be biologically relevant, and should not start with 0. Can be left blank if not provided. |  |
| **Input Protein Length** | Optional | The length (number of positions) of a protein target sequence submitted by a user. | OBI:0002892 | Int | 219 | Report the number of amino acids in the sequence without any units. |  |
| **Input Protein Start** | Optional | The position of the first amino acid in a protein sequence being analyzed (input protein sequence). | OBI:0002975 | Int | 6 | PHA4GE recommends that this value be biologically relevant, and start with 1, however starting at 0 is permissible. Can be left blank if not provided. |  |
| **Input Protein Stop** | Optional | The position of the last amino acid in a protein sequence being analyzed (input protein sequence). | OBI:0002976 | Int | 307 | The value should be biologically relevant, and should not start with 0. Can be left blank if not provided. |  |
| **Input Sequence ID** | Optional | An identifier of molecular sequence or entries from a molecular sequence database. | GENEPIO:0001800 | String | DAAGAT010000041.1 | Can be left blank if not provided. |  |
| **Nucleotide mutation** | Optional | The nucleotide sequence change(s) detected in the sequence being analyzed compared to a reference, represented by a specified notation system. | GENEPIO:0100402 | String | c.1850_1851insGT | Use HGVS syntax; c. (coding) g. (genomic), or r. (ribosomal) mutations should be included here |  |
| **Nucleotide mutation interpretation** | Optional | A description of identified nucleotide mutation(s) that facilitate clinical interpretation. | GENEPIO:0100403 | String | Insertion of GT between nucleotide 1850 and 1851 in katG | Generated computationally from HGVS string |  |
| **Predicted Phenotype** | Optional | A characteristic of an organism that is predicted rather than directly measured or observed. | GENEPIO:0100404 | String | Expected to contribute to antimicrobial resistance | For all AMR tools expects "contributes to resistance" |  |
| **Predicted Phenotype Confidence Level** | Optional | The level of confidence in a predicted phenotype. | GENEPIO:0100405 | String | Low, Medium, High | Confidence that we have that the mutation contributes to the phenotype |  |
| **Amino acid mutation** | Optional | The amino acid sequence change(s) detected in the sequence being analyzed compared to a reference, represented by a specified notation system. | GENEPIO:0100406 | String | p.Ser450Leu | Use HGVS syntax; p. (protein) mutations should be included here; use of 3 letter code preferred for amino acids |  |
| **Amino acid mutation interpretation** | Optional | A description of identified amino acid mutation(s) that facilitate clinical interpretation. | GENEPIO:0100407 | String | amino acid Ser450 is changed to Leucine in rpoB | Generated computationally from HGVS string |  |
| **Reference Gene Length** | Optional | The length (number of positions) of a gene reference sequence retrieved from a database. | OBI:0002888 | Int | 657 | Report the number of nucleotides in the sequence without any units. |  |
| **Reference Gene Start** | Optional | The position of the first nucleotide in a reference gene sequence (sequence being used for comparison). | OBI:0002981 | Int | 1 | The value should be biologically relevant, and should not start with 0. Can be left blank if not provided. |  |
| **Reference Gene Stop** | Optional | The position of the last nucleotide in a reference sequence (sequence being used for comparison). | OBI:0002982 | Int | 900 | The value should be biologically relevant, and should not start with 0. Can be left blank if not provided. |  |
| **Reference Protein Length** | Optional | The length (number of positions) of a protein reference sequence retrieved from a database. | OBI:0002889 | Int | 219 | Report the number of amino acids in the sequence without any units. |  |
| **Reference Protein Start** | Optional | The position of the first amino acid in a reference protein sequence (sequence being used for comparison). | OBI:0002979 | Int | 1 | The value should be biologically relevant, and should not start with 0. Can be left blank if not provided. |  |
| **Reference Protein Stop** | Optional | The position of the last amino acid in a reference protein sequence (sequence being used for comparison). | OBI:0002980 | Int | 300 | The value should be biologically relevant, and should not start with 0. Can be left blank if not provided. |  |
| **Resistance mechanism** | Optional | Antibiotic resistance mechanisms evolve naturally via natural selection through random mutation, but it could also be engineered by applying an evolutionary stress on a population. | ARO:1000002 | String | target alteration | Standardized terms from the ChEBI ontology should be used. Find terms using https://www.ebi.ac.uk/ols/ontologies/aro. Can be left blank if not provided. |  |
| **Strand Orientation** | Optional | The orientation of a genomic element on the double-stranded molecule. | OBI:0002876 | String | + | Values should be sense or antisense, or the corresponding short form "+" or "-". The terms "positive" and "negative" should be avoided. Can be left blank if not provided. |  |

**Supplemental Table 2:** AMR detection tool list.

| Name | Method | Input Type | Description | Mandatory missing hAMRionization fields | Source code |
| --- | --- | --- | --- | --- | --- |
| **Abricate**<br>(Seeman et al., 2015) | BLAST | Contigs | Mass screening of contigs for antimicrobial resistance or virulence genes. It comes bundled with multiple databases: NCBI, CARD, ARG-ANNOT and Resfinder. | analysis_software_version<br>reference_database_version | https://github.com/tseemann/abricate |
| **AMRfinder Plus**<br>(Feldgarden et al., 2021) | BLAST or HMMER | Contigs, ORFs | Find acquired AMR in bacterial protein or assembled nucleotide sequences, as well as known resistance-associated point mutations for several taxa, with NCBI's database. | analysis_software_version<br>reference_database_version<br>input_file_name | https://github.com/ncbi/amr |
| **AMRplusplus**<br>(Lakin et al., 2017) | BWA | Reads | A comprehensive database of antimicrobial resistance genes and user-friendly pipeline for analysis of high-throughput sequencing data | analysis_software_version<br>reference_database_version<br>input_file_name | https://github.com/meglab-metagenomics/amrplusplus_v2 |
| **Ariba**<br>(Hunt et al., 2017) | Read Mapping + Localized Assembly | Reads | Identifies antibiotic resistance genes by running local assemblies. Supports the download of the ARG-ANNOT, CARD, MEGARes, NCBI and Resfinder databases. | analysis_software_version<br>reference_database_name<br>reference_database_version<br>input_file_name | https://github.com/sanger-pathogens/ariba |
| **c-SSTAR (Man & Limbago, 2016)** | BLAST | Contigs | It combines a locally executed BLASTN search against a customizable database with an intuitive graphical user interface for identifying antimicrobial resistance (AR) genes from genomic data. Although the database is initially populated from a public repository of acquired resistance determinants (i.e., ARG-ANNOT), it can be customized for particular pathogen groups and resistance mechanisms. | analysis_software_version<br>reference_database_name<br>reference_database_version<br>input_file_name | https://github.com/tomdeman-bio/Sequence-Search-Tool-for-Antimicrobial-Resistance-SSTAR- |
| **DeepARG**<br>(Arango-Argoty et al., 2018) | DIAMOND + Feed-Forward Neural Network | Reads, Contigs | A deep learning based approach for antibiotic resistance gene annotation. The models were trained from CARD, ARDB and UNIPROT. | analysis_software_version<br>reference_database_version<br>input_file_name | https://github.com/gaarangoa/deeparg2.0 |
| **fARGene**<br>(Berglund et. al., 2019) | HMMER | Reads | A tool that takes either fragmented metagenomic data or longer sequences as input and predicts and delivers full-length antibiotic resistance genes as output. | analysis_software_version<br>reference_database_version<br>input_file_name | https://github.com/fannyhb/fargene |
| **Groot (Rowe & Winn, 2018)** | K-mer + Indexed variant graphs | Reads | Identifies resistance genes from metagenomic samples. As default, it uses the ARG-ANNOT database, with options to use ResFinder, CARD, GROOT-DB and GROOT-CORE-DB. | analysis_software_version<br>reference_database_name<br>reference_database_version<br>input_file_name | https://github.com/will-rowe/groot |
| **KmerResistance (Clausen et al., 2016)** | KMA | Reads | KmerResistance correlates mapped genes with the predicted species of WGS samples, where this allows for identification of genes in samples which have been poorly sequenced or high accuracy predictions for samples with contamination. | analysis_software_version<br>reference_database_version<br>input_file_name | https://bitbucket.org/genomicepidemiology/kmerresistance/src/master/ |
| **Mykrobe (Hunt et al., 2019)** | K-mer + Indexed variant graphs | Reads | Identifies species and resistance profiles for Staphylococcus aureus and Mycobacterium tuberculosis with its own database. |  | https://www.mykrobe.com/ |
| **Pointfinder (Zankari et al., 2017)** | KMA | Reads | The method detects chromosomal mutations predictive of drug resistance based on WGS data. | analysis_software_version<br>reference_database_version<br>input_file_name | https://bitbucket.org/genomicepidemiology/pointfinder/src |
| **Resfams (Gibson et al., 2015)** | HMMER | Contigs | Provides a core database with unique AMR protein sequences from CARD, LacED, and Jacoby and Bush's Beta-lactamase collection. | analysis_software_version<br>reference_database_version<br>input_file_name | https://github.com/dantaslab/resfams |
| **Resfinder (Zankari et al., 2012)** | BLAST or K-mer | Reads, Contigs | ResFinder identifies acquired antimicrobial resistance genes and/or chromosomal mutations in total or partial sequenced isolates of bacteria. | analysis_software_version<br>reference_database_version<br>input_file_name | https://bitbucket.org/genomicepidemiology/resfinder.git |
| **RGI (Alcock et al., 2023)** | BLAST or DIAMOND | Contigs | Predicts resistomes from protein or nucleotide data based on homology and SNP models. It uses reference data from CARD. | analysis_software_version<br>reference_database_version<br>input_file_name | https://github.com/arpcard/rgi |
| **sraX (Panunzi, 2020)** | BLAST or DIAMOND | Contigs | Pipeline to perform resistome analysis in assembled sequences to systematically detect the presence of AMR determinants (ARGs and SNP presence), including the putative detection of new variants. Allows the creation and compilation of local AMR databases. It includes CARD and ARGminer in the package. | analysis_software_version<br>reference_database_name<br>reference_database_version<br>input_file_name | https://github.com/lqpdevtools/sraX |
| **srst2 (Inouye et al., 2014)** | Read Mapping | Reads | This program is designed to take Illumina sequence data, an MLST database and/or a database of gene sequences (e.g. resistance genes, virulence genes, etc) and report the presence of STs and/or reference genes. | analysis_software_version<br>reference_database_version<br>input_file_name | https://github.com/katholt/srst2 |
| **staramr (Bharat et al., 2022)** | BLAST | Contigs | Scans bacterial genome contigs against the ResFinder, PointFinder, and PlasmidFinder databases and compiles a summary report of detected AMR. | analysis_software_version<br>reference_database_version | https://github.com/phac-nml/staramr |
| **TBprofiler (Phelan et al., 2019)** | BWA | Reads | Profiling tool for Mycobacterium tuberculosis to detect resistance and lineage from WGS data |  | https://github.com/jodyphelan/TBProfiler |

